# Stimulus-aware spatial filtering for single-trial neural response and temporal response function estimation in high-density EEG with applications in auditory research

**DOI:** 10.1101/541318

**Authors:** Neetha Das, Jonas Vanthornhout, Tom Francart, Alexander Bertrand

**Affiliations:** KU Leuven, Dept. Electrical Engineering (ESAT), Stadius Center for Dynamical Systems, Signal Processing and Data Analytics. Kasteelpark Arenberg 10, B-3001 Leuven, Belgium; KU Leuven, Dept. Neurosciences, ExpORL. Herestraat 49 bus 721, B-3000 Leuven, Belgium

**Keywords:** EEG processing, temporal response function, speech entrainment, spatial filtering, denoising, forward modeling, attention decoding

## Abstract

A common problem in neural recordings is the low signal-to-noise ratio (SNR), particularly when using non-invasive techniques like magneto- or electroencephalography (M/EEG). To address this problem, experimental designs often include repeated trials, which are then averaged to improve the SNR or to inform the design of a spatial filter that projects the data onto high-SNR directions. However, collecting enough repeated trials is often impractical and even impossible in some paradigms. Therefore, we present a data-driven spatial filter design that takes advantage of the knowledge of the presented stimulus, to achieve a joint noise reduction and dimensionality reduction without the need for repeated trials. The method uses the stimulus-driven neural response, which is then used to find a set of spatial filters that maximize the SNR based on a generalized eigenvalue decomposition. As the method is fully data-driven, the dimensionality reduction enables researchers to perform their analyses without having to rely on their knowledge of brain regions of interest, which increases accuracy and reduces the human factor in the results. In the context of neural tracking of a speech stimulus, our method resulted in better short-term temporal response function (TRF) estimates, higher correlations between predicted and actual neural responses, and higher attention decoding accuracies compared to existing TRF-based decoding methods. We also provide an extensive discussion on the central role played by the generalized eigenvalue decomposition in various denoising methods in the literature, and address the conceptual similarities and differences with our proposed method.

## I. Introduction

Understanding how auditory stimuli influence neural activity is one of the goals of auditory neuroscience research. Towards this goal, several methods have been proposed to model the relationship between the auditory stimuli and the elicited neural responses. In magnetoencephalography (MEG) or electroencephalography (EEG) studies, forward models based on linear temporal response functions (TRFs) are often used to model the path between the stimulus and each of the electrodes/sensors (Ding and Simon, 2012b; Lalor and Foxe, 2010). In this work, we focus on the EEG modality but the methods discussed are not limited to EEG. In case of speech stimuli and multitalker scenarios, TRFs have often been used to linearly map speech stimulus features such as stimulus envelopes, speech spectrograms, phonemes, etc. of both attended and unattended speakers, to the neural activity of the listener (Di Liberto et al., 2015; Ding and Simon, 2012a,b; Power et al., 2012). TRFs not only have high temporal precision but are also sensitive to attentional modulation (Akram et al., 2017; Power et al., 2012). Forward modeling thus comes with the advantages of being able to investigate the TRFs and gain a better understanding of how our brain handles auditory stimuli (Di Liberto and Lalor, 2016; Ding and Simon, 2012a,b), and also the possibility to identify the brain regions involved with stimulus processing (Das et al., 2016; Ding and Simon, 2012b; Etard et al., 2018; Power et al., 2012).

On the other hand, linear backward modeling, where the stimulus features are reconstructed from the neural activity, is also a commonly used method (Biesmans et al., 2017; Ding and Simon, 2012b; Fuglsang et al., 2017; Mesgarani and Chang, 2012; Mirkovic et al., 2015; O’Sullivan et al., 2014). Unlike the forward model, this approach makes use of inter-channel covariances to design the decoder. The correlations between the reconstructed and the original stimulus features are thus higher than those of the predicted and original neural responses in the forward modeling approach. However, the decoder coefficients themselves can not directly be interpreted, unlike forward models, where TRFs for different channels can be visualized using topoplots.

In algorithms that deal with neural activity, often dimensionality reduction is key. Dimensionality reduction works on the assumption that the data of interest lies in a lower dimensional space than its original representation. As mentioned earlier, forward models map the auditory stimulus to each of the electrodes, thus preserving spatial information of stimulus-related cortical activity. However, unlike algorithms that employ backward modeling, they do not use cross-channel information to regress out non-stimulus related activity (Wong et al., 2018). In such cases, dimensionality reduction can help to transform neural recordings from a multi-electrode system, into a signal subspace with fewer components (than electrodes) and better SNR of the stimulus following responses (Akram et al., 2016, 2017; Ding et al., 2014). The algorithm may then use these components themselves to achieve its goal, or project the components back to the electrode space, effectively performing a denoising operation, before further processing. Thus, dimensionality reduction goes hand-in-hand with denoising.

Principal component analysis (PCA) is often used for dimensionality reduction, in which case the principal components corresponding to lower variance are discarded (Cunningham and Byron, 2014; de Cheveigné et al., 2018b). However, this approach relies on the assumption that low variance corresponds to non-relevant activity, which can be a rather restrictive assumption to make, particularly for EEG data, where usually most of the variance comes from artifacts and background neural activity that is unrelated to the stimulus. Another approach, independent component analysis (ICA) (Bell and Sejnowski, 1995), works on the assumption that the components (or sources) are statistically independent. It is often used to remove components corresponding to artifacts with specific patterns (like eyeblinks) (Urigüen and Garcia-Zapirain, 2015). However, ICA does not perform well in the extraction of signal components that are far below the noise floor, as is the case for neural responses to speech. Another method, joint decorrelation (JD) (de Cheveigné and Parra, 2014) follows the formulation of linear denoising source separation (DSS) (Särelä and Valpola, 2005) to improve the SNR of the activity of interest in the neural data. In the most commonly used version of DSS, a criterion of stimulus-evoked reproducibility is used, maximizing the evoked-to-induced ratio. This, however, requires repeated trials, which renders it impractical for many EEG applications (although for MEG data, a few trials are typically enough for DSS to obtain a useful dimensionality reduction and denoising (Akram et al., 2016, 2017; Ding et al., 2014)). Canonical correlation analysis (CCA) also reduces dimensionality by finding separate linear transformations for the stimulus as well as neural responses, such that in the respective projected subspaces, the neural response and the stimulus are maximally correlated (de Cheveigné et al., 2018a,b; Dmochowski et al., 2018; Hotelling, 1936).

Except for CCA, the methods mentioned above do not exploit the knowledge of the stimulus. The goal of this work was to develop a joint dimensionality reduction and denoising algorithm for neural data which takes advantage of the knowledge of presented stimulus. We propose such a data-driven stimulus-aware method, which finds a set of spatial filters using the generalized eigenvalue decomposition, to maximize the SNR of stimulus following responses, thereby also facilitating an accurate TRF estimation on shorter trial lengths. For neural responses to continuous speech stimuli, we show that the proposed method results in an effective dimensionality reduction/denoising without the need for data from repeated stimulus trials as in the DSS method or phase-locked averaging techniques.

We validate the performance of the proposed method in the following three contexts.

1. Short-term TRF estimation, where short trials are used to estimate TRFs that map the auditory stimulus envelope to the neural responses. The estimated TRFs can be visualized to track the effect of attention on the TRF shapes, and eventually to even decode attention in real-time without any prior training of decoders (Akram et al., 2017; Miran et al., 2018).
2. Stimulus envelope tracking, where in a single speaker scenario, neural responses to the stimulus envelope are predicted using forward modeling, i.e., finding TRFs that map the stimulus envelope to the neural responses. The correlations between the original neural responses and the predicted stimulus following responses are analyzed (Aiken and Picton, 2008; Di Liberto et al., 2015). The analysis of these correlations can not only contribute to advancing our knowledge of how the brain responds to auditory stimuli under different conditions, but also has potential to act as objective measures of speech intelligibility (Broderick et al., 2018; Di Liberto et al., 2018; Ding and Simon, 2012a; Lesenfants et al., 2019; Vanthornhout et al., 2018).
3. Auditory attention decoding, where an estimate is made as to which of multiple speakers, a person is attending, using forward modeling (Wong et al., 2018). Attention decoding finds applications in brain computer interfaces (BCIs) such as neuro-steered hearing prostheses where accurate information about a person’s auditory attention can be used to steer the noise suppression beamformer towards the direction of the attended speaker (Aroudi et al., 2018; Das et al., 2017; O’Sullivan et al., 2017; Van Eyndhoven et al., 2017). In particular, for real-time tracking of auditory attention, forward models are popular (Akram et al., 2016, 2017; Miran et al., 2018), in which case, dimensionality reduction is an important ingredient.

We compare our method with the commonly used DSS method, which is known to outperform PCA and ICA for estimation of weak neural responses, such as those obtained from auditory stimuli (de Cheveigné and Simon, 2008).

Furthermore, we show a non-straightforward algebraic link between our method and the CCA method, showing that CCA can be viewed as a special case of the presented method. A direct consequence is that CCA can also be used in a context of EEG denoising or short-term TRF estimation.

The outline of the paper is as follows. In Section II, we describe the algorithm for denoising and dimensionality reduction. In Section III, we describe the validation within the three aforementioned contexts. The strengths of our method as well as similarities and differences with existing denoising and dimensionality reduction methods are discussed in Section IV. Finally, we summarize and conclude in Section V.

## II. Algorithm

The main goal of this work is to find a spatial filter, or a group of spatial filters, that combine the EEG channels in such a way that the power of the auditory stimulus following responses is increased in comparison to the rest of the neural activity captured by the electrodes. We aim to achieve this goal using the following steps to train the filters: 1) Estimate the spatial covariance of the desired neural response across the different EEG channels, using the known stimulus as side information, 2) find a signal subspace within which the ratio of the power of the desired neural response to the power of the background EEG signal (i.e. SNR) is maximized, 3) project the EEG data into the new signal subspace, thereby performing a joint dimensionality reduction and denoising, 4) back-project the denoised data into the channel space if necessary. After step 3 or 4, one can then use the denoised signals to perform the desired task-related analysis such as, e.g., estimating per-trial TRFs or correlating the stimulus with the denoised EEG data to more accurately quantify the neural entrainment. The steps of the algorithm are illustrated in figure 1 and explained in detail below. Note that the split in training and testing data as indicated in figure 1 was done for validation purposes.

**Fig. 1:**
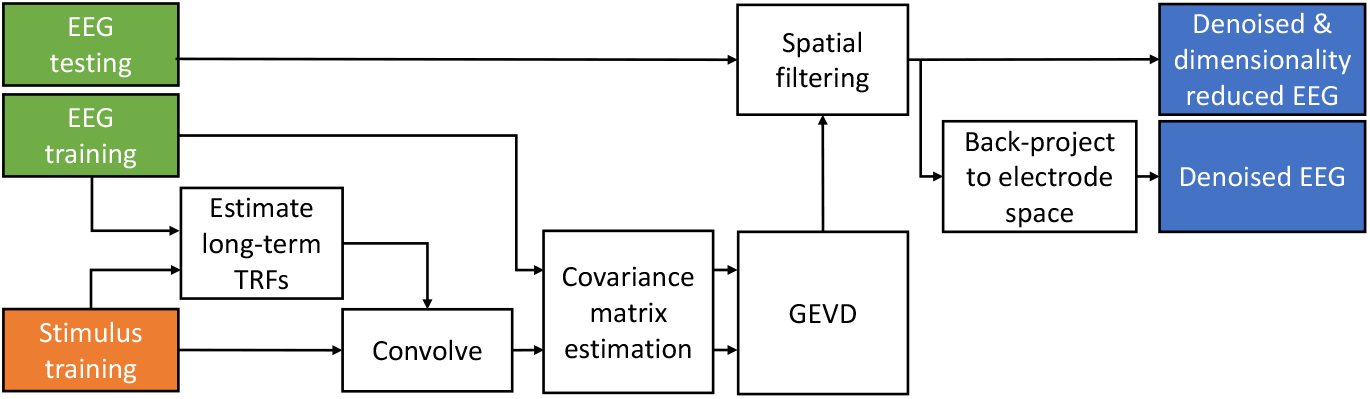
Block diagram of the proposed method. The EEG and stimulus data in the training set is used for forward modeling. The resulting TRFs are convolved with the stimulus to form stimulus following responses (desired signal). A GEVD on the covariance matrices of these signals and that of the original EEG results in the spatial filter that can be used to improve the SNR of the test EEG data. Note that the split in training and testing data was done for validation purposes.

### A. Max-SNR formulation

The EEG signal, at a time index *t*, is defined as a *C*-dimensional vector **m**(*t*) = [*m*_1_(*t*), *m*_2_(*t*), …*m_C_*(*t*)]^*T*^ ∈ ℝ^*C*^ where *C* denotes the number of channels, and *m_i_*(*t*) represents the EEG sample from the *i^th^* channel at time index *t*. For example, in a scenario where the subject listens to a speech signal *s*(*t*), it is known that the neural responses entrain to the envelope of the speech stream. Therefore, the EEG signal can be assumed to be the sum of the activity driven by the speech stimulus **x**(*t*) and the rest of the neural response **n**(*t*) at time *t*.

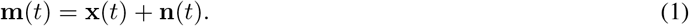

Our goal is to find a set of spatial filters that maximizes the SNR at their outputs. Consider the matrix **P**_*K*_ ∈ ℝ^*C*×*K*^ which contains **K** spatial filters in its columns. By multiplying the EEG data **m**(*t*) with 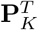, the *C* EEG channels are combined into *K* output channels where *K* < *C*, in such a way that the SNR at the output is maximized. For the sake of simplicity, we first assume *K* = 1, thus reducing **P**_*K*_ to a single spatial filter denoted by the vector **p** ∈ ℝ^*C*^. The maximum-SNR criterion thus becomes:

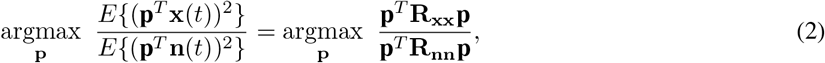

Assuming the stimulus following responses **x**(*t*) and the background EEG (noise) **n**(*t*) are uncorrelated, the maximal SNR criterion is equivalent to solving the optimization problem of maximizing the signal to signal plus noise ratio (SSNR) (see Appendix A for the proof) such that equation (2) is equivalent to the new equation

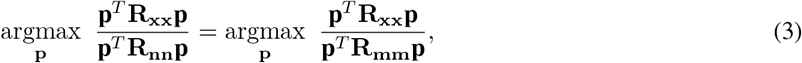

where **R_mm_** = *E*{**m**(*t*)**m**(*t*)^*T*^} ∈ ℝ^*C×C*^ (mean centered) contains both noise and desired neural responses. Thus the max-**SNR** formulation requires two covariance matrices – **R_xx_** of the stimulus-following responses, and **R_mm_** of the original EEG data. **R_mm_** can easily be computed from the raw EEG data, while the estimation of **R_xx_** is explained in the next subsection.

### B. Stimulus-following neural response covariance estimation

The stimulus following neural response **x**(*t*) can be modeled by a linear temporal response function (TRF) (Ding and Simon, 2012a,b; Lalor and Foxe, 2010) that maps the stimulus envelope (including *N_ι_*-1 time lagged versions of it) *s*(*t*) = [*s*(*t*), *s*(*t* – 1),…, *s*(*t* – *N_ι_* + 1)]^*T*^ ∈ ℝ^N_ι_^ to the neural response:

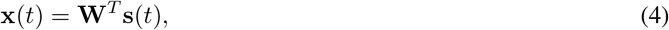

where **W** ∈ ℝ^*N_ι_*×*C*^ is a matrix containing the per-channel TRFs in its columns. **W** can be estimated by minimizing the mean square error between **x**(*t*) and **m**(*t*):

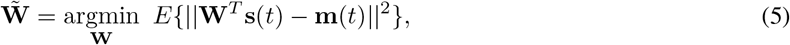

where *E*{.} denotes the expected value operator. The solution of (5) is given by

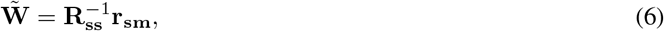

where **R_ss_** = *E* {**s**(*t*)*s*(*t*)^*T*^} ∈ ℝ^*N_ι_ ×N_ι_*^ is the covariance matrix of the stimulus envelope, and **r_sm_** = *E* {**s**(*t*)**m**(*t*)^*T*^} ∈ ℝ^*N_ι_×C*^ is the cross-correlation matrix between the stimulus envelope and the EEG data.

Consider we have *N* samples of data, with which to estimate the TRF 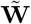. By concatenating the **N** vectors **s**(*t*) and **m**(*t*) in the columns of the matrices **S** = [**s**(1),**s**(2),…,**s**(*N*)] ∈ ℝ^*N*_*ι*_×*N*^ and **M** = [**m**(1), **m**(2),…, **m**(*N*)] ∈ ℝ^*C×N*^, respectively, we can estimate the covariance matrix of the stimulus envelope as **R_ss_** ≈ (**SS**^*T*^)/*N* and the cross-correlation matrix between the stimulus envelope and the EEG data as **R_sm_** ≈ (**SM**^*T*^)/*N* which can then be used in (6) to estimate the TRF matrix 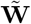 as

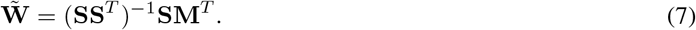

With the estimated TRF, the desired neural response **X** = [**x**(1), **x**(2),…, **x**(*N*)] ∈ ℝ^*C×N*^ can be computed using (4), the spatial covariance matrix of which can then be estimated as **R_xx_** ≈ (**XX**^*T*^)/*N*.

Besides knowledge of the stimulus itself, the algorithm can also be used to take advantage of the knowledge of the dependence of the TRF with respect to various conditions. For example, it is already known that the TRFs for speech stimuli from the right side of the listener have different spatial patterns compared to TRFs for speech stimuli from the left side of the listener (Das et al., 2016; Power et al., 2012). One way of exploiting this information would be by separating the data into ‘attention to the left’ and ‘attention to the right’ conditions, and separately finding long-term TRFs **W_L_** and **W_R_** respectively. The desired neural responses **x_L_**(*t*) and **x_R_**(*t*), and consequently the corresponding spatial covariance matrices **R_xxL_** and **R_xxR_** can then be estimated separately. In this manner, we estimate multiple spatial covariance matrices that collectively model the condition-dependencies in the second order statistics of the desired neural responses. In this case, we can maximize the *average* SNR across both conditions:

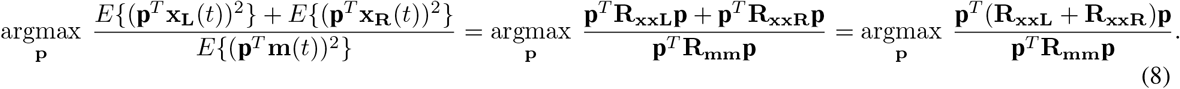

As a result, the matrix **R_xx_** in equation (3) is replaced with **R_xxL_** + **R_xxR_**. We will demonstrate that this additional modeling freedom can be exploited to design better spatial filters. Note that a more naive model in which a single TRF is fitted over all conditions would result in a virtual TRF that does not match any of the true underlying response functions, leading to a single **R_xx_** matrix which is *not* equal to **R_xxL_** + **R_xxR_**.

### C. Spatial filter estimation

Having estimated the spatial covariance matrix **R_xx_** of the desired neural response **x**(*t*) (or an averaged **R_xx_** across multiple conditions as in equation (8)), the next step is to find the set of spatial filters that maximizes the SNR at their outputs. Solving for the max-SNR criterion, the stationary points of equation (3) can be shown to satisfy (Van Veen and Buckley, 1988)

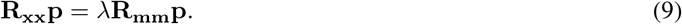

which defines a generalized eigenvalue problem for the matrix pencil (**R_xx_**, **R_mm_**) (Biesmans et al., 2015; Golub and Van Loan, 1996; Parra and Sajda, 2003), where all λ’s and p’s that satisfy equation (9) are denoted as the generalized eigenvalues and eigenvectors, respectively. Note that equation (9) can be transformed in an equivalent eigenvalue problem

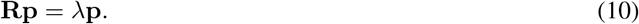

involving the non-symmetric matrix 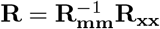. From equation (9), it follows that

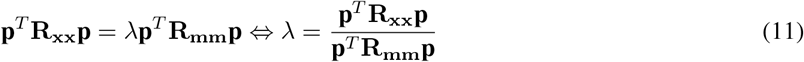

which implies that λ is equal to the output SNR of the spatial filter. Therefore, in order to maximize the SNR, we should set **p** equal to the (generalized) eigenvector corresponding to the largest (generalized) eigenvalue λ.

So far, we have considered a spatial filter **p** which combines the *C* EEG channels in an optimal way to obtain a channel with SNR maximized. This can be further extended to solving the problem of finding a filter bank **P**_*K*_ ∈ ℝ^*C×K*^ consisting of *K* spatial filters that maximizes the total output SNR, by finding *K* generalized eigenvectors corresponding to the *K* highest eigenvalues from the GEVD of (**R_xx_**, **R_mm_**). **R_mm_** can be computed from the raw EEG data as **R_mm_** ≈ (**MM**^*T*^)/*N* where **M** is zero-centered. We refer to the resulting spatial filterbank as the stimulus-informed GEVD (SI-GEVD) filter.

Although they were originally described as a sequence of two PCA steps, algorithms like JD and DSS can also be realized as a single decomposition by means of a GEVD of two covariance matrices. The key difference with our approach is that in DSS and JD, the **R_xx_** matrix is replaced with a surrogate covariance matrix in which the contribution of the stimulus following response is enhanced with respect to the background EEG. This can be achieved through averaging the neural responses over multiple repetition trials, or using application specific filters or selecting high SNR epochs. However, since the resulting ‘enhanced’ covariance matrix still contains a residual covariance of the (distorted) background EEG, the resulting filters will not be SNR-optimal. Furthermore, if this residue becomes very high, e.g., because insufficient repeated trials are available for averaging, the filters provided by DSS/JD method will not be of any practical value. Instead, in the SI-GEVD method, we avoid this averaging step with a forward convolutional model between the continuous stimulus and the different EEG channels in order to estimate the spatial second-order statistics of the neural response.

As mentioned in the introduction, the CCA method can also be viewed as a stimulus-aware subspace method. However, as opposed to SI-GEVD, CCA manipulates the stimulus and the EEG data simultaneously to find components on both sides with a maximal mutual correlation. Because of this joint manipulation with correlation maximization as the main target, CCA is at first sight only remotely related to max-SNR approaches like SI-GEVD, JD, or DSS, as its goal is not to maximize SNR directly. Remarkably, it can be shown that CCA is actually a special case of SI-GEVD, i.e., when it is used in its simplest form with a single TRF estimate as in equation (3) (we provide a proof in Appendix B). This also shows that there is a link between DSS, JD and CCA, where the ‘glue’ between these methods is provided by the SI-GEVD formulation. The equivalence is not obvious at first sight and only holds if the filters applied on the EEG side are purely spatial and do not include temporal statistics (as opposed to the stimulus-side, where also time lags can be used). As a direct consequence, CCA can also be used as a denoising algorithm, e.g., to estimate short-term TRFs as explained in the next section. However, the SI-GEVD has a more general model and allows for more flexibility, e.g., to incorporate condition-dependent variability in the TRFs, such as the example given in equation (8). If we take into consideration the direction of the stimulus, the optimization problem can be defined in such a way that the average SNR over all conditions (in our earlier example, ‘attention to the left’ and ‘attention to the right’) is maximized, as formulated in equation (8). The SI-GEVD filter can then be found from the GEVD of the matrix pencil (**R_xxL_** + **R_xxR_**, **R_mm_**). This will lead to a bank of spatial filters which aims to find a subspace that leads to a high SNR in *both* of these conditions. Note that, while the training phase requires side information about the direction of attention, this information is not required during the test phase after the spatial filters have been trained.

### D. Project EEG signal onto the new subspace

The top *K* eigenvectors from the GEVD corresponding to the *K* largest generalized eigenvalues λ can be used to reduce the dimensionality of the raw EEG data with *C* channels, to *K* filtered EEG components. If **P**_*K*_ = [**p**_1_, **p**_2_,…, **p**_*K*_] is the matrix containing the top *K* eigenvectors in its columns, then the compressed EEG signal in the new subspace is

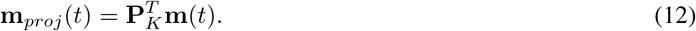

There are many ways to choose *K*. The cumulative λ values can be plotted and the top *K* values which contribute to 95% of the total SNR (sum of all λ values) can be chosen. The optimum *K* can also be chosen based on the application. For example, in the case of auditory attention decoding using forward modeling or correlation analysis, one can use cross validation to select the number of components *K* that eventually results in the highest attention decoding accuracy or the best correlations respectively.

### E. Back-projection to the electrode space

If desired^1^, back-projection to the electrode space can be done based on the matrix 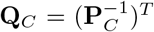, where the square matrix **P**_*C*_ contains all the generalized eigenvectors of the pencil (**R_xx_**, **R_mm_**) in its columns. Note that **P**_*K*_ contains a subset of the columns of **P**_*C*_. By selecting the subset with the same column indices from **Q**_*C*_, we obtain the matrix **Q**_*K*_, which can be used to project the compressed data **m**_*proj*_(*t*) back to the electrode space with minimal error in the least squares sense (see Appendix C):

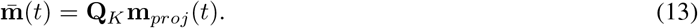

This results in a denoised version 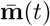 of the original EEG channels **m**(*t*). The denoised signals can then be used for short-term TRF estimation, attention decoding, etc.

## III. Validation Experiments

### A. Data collection and pre-processing

#### 1) Dataset I

This dataset consists of EEG data collected during the experiments described in Das et al. (2016). The data consists of 64-channel EEG recordings sampled at 8192 Hz collected from 16 normal hearing subjects (8 male, 8 female) between the age of 17 and 30 years. The subjects were asked to attend to one of two stories that were simultaneously presented to them. We refer to Das et al. (2016) for details on the experiment conditions. For the analyses in this paper (including those in section III-D), we used only the data corresponding to conditions where the presented stimuli were filtered by head-related transfer functions (HRTFs) (≈38 minutes per subject), in order to provide a realistic acoustic impression of the location of the speakers. In Das et al. (2016), it has been shown that this data resulted in significantly better attention decoding performance in comparison to dichotically presented unfiltered stimuli. The multi-channel Wiener filtering (MWF) method in Somers et al. (2018) was used on the EEG data for artifact removal. The EEG data was bandpass filtered between 1 and 9 Hz. In Das et al. (2016); Ding and Simon (2012b); Golumbic et al. (2013); Pasley et al. (2012), it was shown that cortical envelope tracking in this frequency range results in the best attention decoding performance. The audio envelope was determined by filtering the speech waveform with a gammatone filterbank (with 15 filters) followed by powerlaw compression (exponent = 0.6) on the absolute value of each filter’s output signal (Biesmans et al., 2017). The resulting signals from all subbands were then summed, after which the signal was downsampled to 32 Hz and filtered using the same 1-9 Hz bandpass filter as for the EEG signal, which then resulted in a smooth envelope. We decided to keep the preprocessing identical to the original paper Das et al. (2016) as it is application specific, and hence has differences with the preprocessing done in section III-A2 (based on Vanthornhout et al. (2018)). The EEG data was referenced to the Cz electrode^2^ (therefore, *C* = 63). All data were normalized to have zero mean and unit standard deviation per channel/lag.

#### 2) Dataset II

This dataset consists of EEG data from Vanthornhout et al. (2018). In this study the EEG (64 channels sampled at 8192 Hz) was recorded from 27 normal hearing subjects (8 male, 19 female, average age: 23 years). The participants listened to a 14-minute continuous speech stimulus (the story Milan, narrated in Flemish by Stijn Vranken) and to isolated sentences. The isolated sentences stimulus consists of a concatenation of 40 sentences, with a 1 s silence between each sentence, taken from the Flemish Matrix sentence test (Luts et al., 2014). As each sentence is approximately 2 s long, this yields a stimulus of 120 s with 80 s of speech. Each isolated sentences stimulus was repeated 3 or 4 times, yielding 6 to 8 minutes of EEG recordings. All stimuli were presented diotically. In the original study (Vanthornhout et al., 2018) the isolated sentences were presented in silence and in noise. For this study, we only used the data without background noise.

Preprocessing of the EEG data was done similar to Vanthornhout et al. (2018). The EEG was highpass filtered (second order Butterworth with cut-off at 0.5 Hz) and downsampled to 256 Hz before applying the MWF for artifact rejection (Somers et al., 2018). It was then re-referenced to the Cz electrode (therefore *C* = 63). Next, the EEG was bandpass filtered within the delta band (0.5 – 4 Hz). This band was found to be the most useful to predict speech intelligibility (Vanthornhout et al., 2018). Finally, the EEG was downsampled to 128 Hz. The stimulus envelope was estimated using a gammatone filterbank with 28 filters. A powerlaw compression was then applied (exponent = 0.6) to the absolute values in each of the subbands. The resulting 28 subband envelopes were bandpass filtered (as was done for the EEG data) and averaged to obtain one single envelope, and downsampled to 128 Hz. The EEG data and stimulus data were normalized (per channel/lag) to have zero mean and unit standard deviation.

#### 3) Dataset III (synthetic dataset)

For this dataset, the EEG data synthesis was based on a real EEG recording from Dataset I, where the ‘mTRF’ toolbox (Crosse et al., 2016) was used to find the long-term TRFs **W_L_** and **W_R_** for the 2 conditions:’attention to the right’ and ‘attention to the left’ separately, mapping the attended speech envelope (and it’s lagged versions up to 400 ms) to the EEG data across all subjects. The lag at which **W_L_** and **W_R_** jointly had the maximum norm was chosen as the reference lag. The left and right attended TRFs corresponding to the channel ‘C5’ was then taken to generate base TRF templates. For each condition, TRFs for all other channels were taken to be scaled versions of this base template, with the scaling corresponding to the amplitude of the TRFs in the corresponding channels at the reference lag. Thus 2 C-channel TRF matrices **W**_baseL_ and **W**_baseR_ ∈ ℝ^*N_ι_*×*C*^ containing the per-channel TRFs in its columns was synthesized. Lags of 0 to 400 ms were used corresponding to *N_ι_* = 32 Hz × 400 ms + 1 = 14 samples.

The stimulus following responses were then synthesized by convolving the per-channel TRFs **W**_baseL_ and **W**_baseR_ with the attended speech envelope and the results were put in matrices **X_L_** and **X_R_** ∈ ℝ^*N×C*^ where *N* is the total number of EEG samples (38 minutes) simulated per condition. Noise **N** to be added to the stimulus following responses **X_L_** and **X_R_** was synthesized by flipping in time, the concatenated 38 minute EEG from a randomly chosen subject. The flipping operation ensures that the neural response in the EEG data is uncorrelated to the synthesized neural response. EEG responses to the attended speech **M** ∈ ℝ^2*N×C*^, which included both the conditions, were simulated by adding noise **N** to **X_L_** and **X_R_** at different SNRs. SNR is defined here as 20 times the logarithm (base 10) of the ratio of the root mean square (rms) values (averaged across channels) of the neural response (**X_L_** or **X_R_**) to the rms values (averaged across channels) of the noise (**N**).

### B. Short-term TRF estimation

We look at the problem of estimating TRFs from short trials. This is different from TRF estimation within the SI-GEVD method itself where we are estimating long-term TRFs over a large amount of data (i.e., N is very large) ensuring goog estimates. Unlike long-term TRF estimation, estimating TRFs from short trials is a challenging task owing to the low SNR of the EEG data and the relatively few number of samples to estimate the TRFs from. Therefore, by SI-GEVD filtering we expect to improve the SNR of the EEG data enabling us to better estimate short-term TRFs over much shorter windows with fewer samples.

#### 1) Data Analysis

In order to reliably estimate any improvement that SI-GEVD filtering might bring to the problem of TRF estimation from short trials, we also need ground-truth data of which the true underlying TRF is exactly known. To this end, we generated hybrid EEG data which consisted of real EEG to which a simulated neural response was added at various possible SNRs, as described in the synthesis of Dataset III. Once the EEG data was synthesized using the two base TRF templates, we focused on estimating short-term TRFs from the SI-GEVD filtered version, and assessing the quality of these TRF estimates in comparison to the base TRF template. We estimated TRFs **W**_raw_ ∈ ℝ^*N_ι_×C*^ from 120 s trials of the unfiltered EEG data M using a standard ridge regression technique as implemented in the mTRF toolbox (Crosse et al., 2016). The regularization parameter for ridge regularization was chosen to be 0.2 after analyzing the results over a range of values. In addition, a leave-one-trial-out cross validation was used for estimating the SI-GEVD filter. This is explained as follows. For the *i*^th^ trial, the EEG data **M**_*test*_[*i*] = [**m**((*i* – 1)*L* + 1), **m**((*i* – 1)*L* + 2),…, **m**(*iL*)] ∈ ℝ^*C×L*^ formed the *i^th^* test trial, while the EEG data from all trials except the test trial, formed the training set **M**_*train*_[*i*] ∈ ℝ^*C*×(2*N–L*)^, where *L* = 32 Hz × 120 s = 3840 samples and *N* = 32 Hz × 120 s × 19 trials per condition = 72960 samples. Similarly, the stimulus envelope of the test trial was taken as **S**_*test*_[*i*] = [**s**((*i* – 1)*L* +1), **s**((*i* – 1)*L* + 2),…, **s**(*iL*)] ∈ ℝ^*N_ι_×L*^ where **s**(*t*) is a vector containing *N_ι_* = 14 samples corresponding to *N_ι_* delays, as seen in equation (4). Stimulus envelopes from all trials except the test trial of index *i*, were concatenated to form the training set **S**_*train*_[*i*] ∈ ℝ^*N_ι_*×(2*N – L*)^.

We estimate the SI-GEVD filters **P**_*K*_ [*i*] on the training data **S**_*train*_ [*i*] and **M**_*train*_ [*i*] based on the method described in section II, taking advantage of the direction of the stimulus (solving equation (8)). Since, in all cases, there were 2 dominant generalized eigenvalues, only 2 generalized eigenvectors (*K* = 2) were used as the SI-GEVD filter. The test trial was then SI-GEVD filtered and back-projected to obtain the denoised EEG data 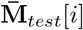. The TRFs **W**_SI-GEVD_[*i*] were then estimated from 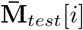 and **S**_*test*_[*i*] within the 120 s trial, in the same way as how **W**_raw_[*i*] were estimated.

As explained in section II-C, CCA can be viewed as a special case of SI-GEVD, which implies it can also be used as a denoising filter, albeit without the flexibility in using side information about different conditions such as left and right-attended. To demonstrate the benefit of this extra modeling power, we also implemented CCA and used it as a denoising filter (similar to SI-GEVD), where the same leave-one-trial-out procedure was used. The back-projection to the electrode space is here performed with a standard least-squares regression, as the equation (13) only works for the generalized eigenvector formulation.

#### 2) Results

For each 120 s trial, the estimated TRFs of all channels were concatenated into a single vector estimated TRF: **ŵ**_raw_[*i*] ∈ ℝ^*CN_ι_*×1^ from the raw EEG data, **ŵ**_CCA_[*i*] ∈ ℝ^*CN_ι_*×1^ from the CCA-filtered data, and **ŵ**_SI-GEVD_ [*i*] ∈ ℝ^*CN_ι_* ×1^ from the SI-GEVD filtered EEG data. For each trial, the base TRFs of all channels (depending on which condition the trial belonged to, i.e., left or right attended) were also concatenated to obtain a single vector base TRF **ŵ**_base_ [*i*] ∈ ℝ^*CN_ι_* ×1^. We omit the indication for the left or right attended stimulus conditions for notational convenience. In order to eliminate any differences due to scaling, a scaling factor was estimated by applying a least squares fitting such that the estimated TRF vector (referred to, in general, as **ŵ**) was scaled to fit **ŵ**_base_ [*i*] in the minimum mean squared error sense. This allows for a more fair comparison, as it allows to compensate for amplitude bias in the back-projection^3^, and since we are mainly interested in the shape and *relative* amplitude of the TRFs across channels. The scaling was performed across channels (note that w is a vector in which the TRFs of all channels are concatenated), and thus the relative differences in the per-channel amplitudes remained unchanged. This scaling factor was found as follows:

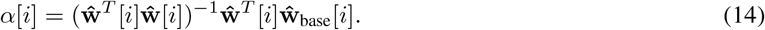

The mean squared error (MSE) between the scaled estimated TRF vector and the base TRF vector was computed for all the trials. The MSE values were normalized by the square of the norm of the base TRF vectors to find relative MSEs in the range of 0 to 1 resulting in the following definition:

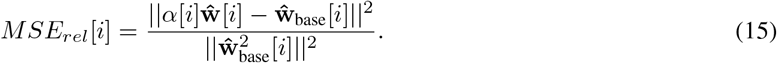

For a range of SNRs, figure 2 shows the relative MSE for TRF estimation from raw EEG, CCA-filtered EEG and SI-GEVD filtered EEG. A typical example of how the different methods perform is demonstrated in figure 3 showing the estimated TRFs compared to the base TRF of channel ‘C5’ for a randomly chosen trial. The relative MSEs for the three approaches were compared using Wilcoxon’s signed-rank test (with Bonferroni correction). For all the SNRs analyzed (0 dB to −25 dB), the relative MSEs from SI-GEVD filtering was found to be significantly lower (*p* < 0.0001) than those from raw data, as well as CCA-filtered data, showing that SI-GEVD filter is effective in denoising the data which translates into better TRF estimates over short trials.

**Fig. 2:**
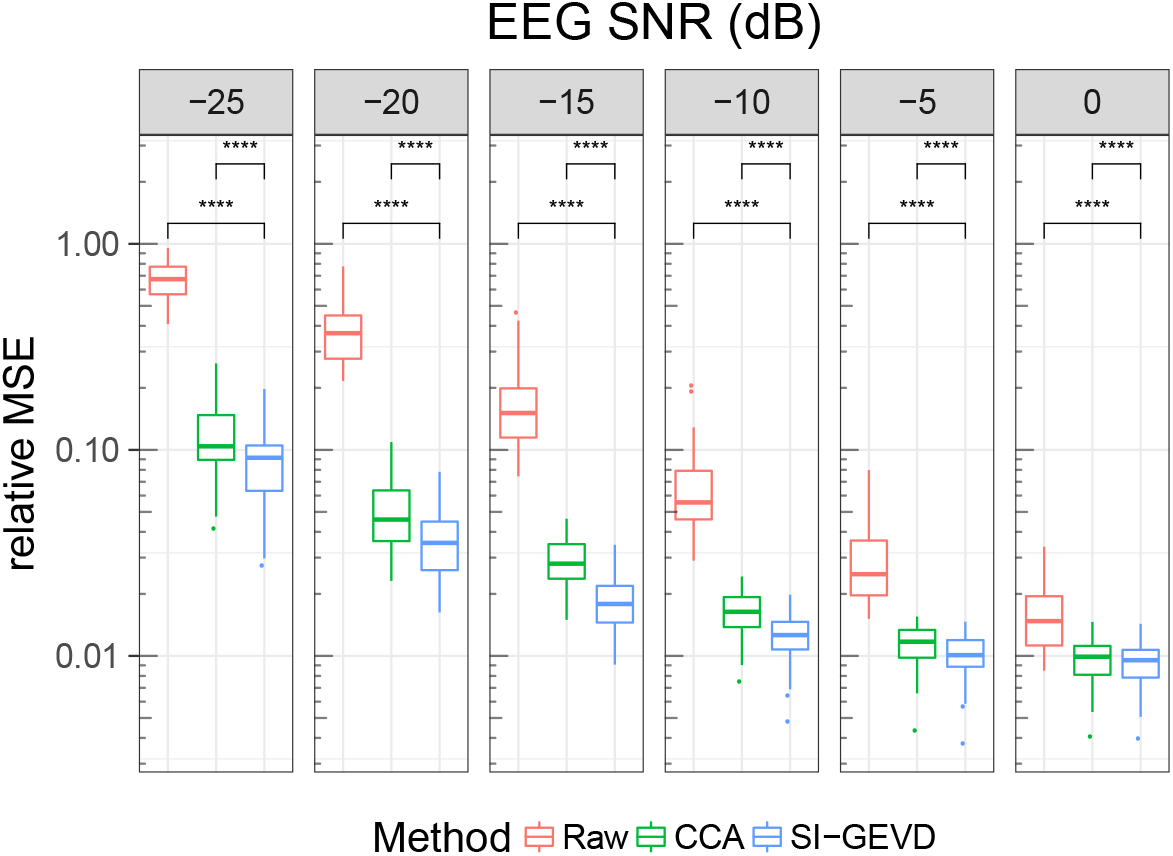
Relative MSE between base TRFs and estimated TRFs based on raw Vs CCA-filtered Vs SI-GEVD filtered EEG. Each boxplot is built from the statistics of per trial relative MSEs (38 datapoints per method). Comparison between the methods were done using Wilcoxon’s signed-rank test with Bonferroni correction: ‘****’ for *p* < 0.0001. Note that the Y-axis is in log scale.

**Fig. 3:**
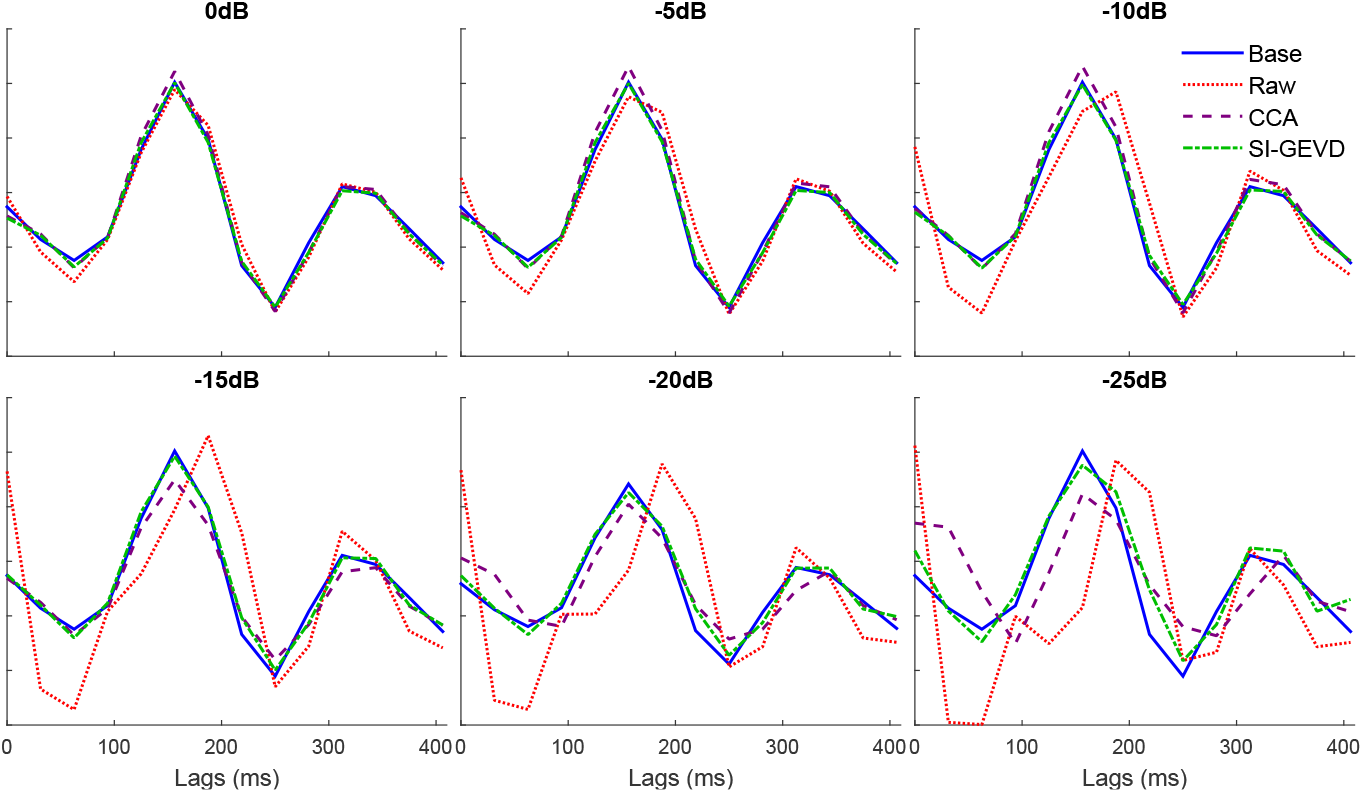
An example base TRF (template) of channel ‘C5’ is compared with the TRF estimated from raw data Vs CCA-filtered data Vs SI-GEVD-filtered data. The plotted TRFs are from a randomly chosen 120 s trial taken from Dataset III (synthetic EEG data).

### C. Speech envelope tracking

The extent of tracking of an auditory stimulus by the neural responses can be analyzed by looking at the correlations between the original EEG data and the predicted stimulus-following responses (estimated by convolving the stimulus envelope with estimated TRFs). Such an analysis can help us understand how the brain responds to auditory stimuli as well as provide objective measures of speech intelligibility. SI-GEVD filtering may be used to boost the SNR of the EEG data which can then result in better correlations between predicted stimulus-following data and the original EEG data.

#### 1) Data Analysis

To analyze the effectiveness of the SI-GEVD filter in measures of envelope tracking, we used the data from Dataset II. Details of the data collection and preprocessing can be found in section III-A2. The goal of this analysis was to use forward modeling to predict speech following neural responses. Correlations of the original EEG and the predicted EEG were then used to quantify the effectiveness of denoising using the SI-GEVD filter. The data from the continuous speech stimulus was used only for training the TRFs, and not for training SI-GEVD filters, or for testing. On the other hand, the isolated sentences was used both in the training as well as the testing set for the estimation of the SI-GEVD filters and the TRFs, while performing leave-one-trial-out cross validation. This dataset was split into 6 trials, each trial consisting of 20 s of EEG data. Thus, for each test trial (isolated sentences from Dataset II), the SI-GEVD filter was trained on the remaining 5 isolated sentences trials and their repetitions (3 repetitions × 5 trials × 20 s), and then the TRFs were trained on a) the remaining 5 isolated sentences trials and their repetitions (3 repetitions × 5 trials × 20 s), and b) the data from the continuous speech stimulus (14 minutes). SI-GEVD filtering can be done on continuous data, the splitting into trials was only done to facilitate cross-validation.

The *i^th^* test trial **M**_*test*_[*i*] ∈ ℝ^*C×L*^ contained *L* = 128 Hz × 20 s = 2560 time samples per channel, while the training set **M**_*train*_[*i*] ∈ ℝ^*C×N*^ consisted of *N* = (3 repetitions of isolated sentences × 5 trials × *L*) + (14 minutes of continuous speech × 60 **s** × 128 Hz) time samples per channel. Stimulus envelopes **S**_*test*_[*i*] ∈ ℝ^*N_ι_×L*^ and **S**_*train*_[*i*] ∈ ℝ^*N_ι_×N*^ consisted of lags up to 75 ms, and hence *N_ι_* = 128 Hz × 75 ms + 1 = 11 samples.

For each trial, the training set data was used to find the SI-GEVD spatial filter **P**_*C*_ [*i*] and the back-projection matrix **Q**_*C*_ [*i*]. After choosing the number of components to be used (the method of choosing *K* is explained further on), the EEG data from each test trial was denoised by applying the SI-GEVD spatial filter found from the corresponding training set, and back-projected to the C-channel space. In this manner, all the EEG data were denoised, and then used for TRF estimation. From the denoised training set, the TRFs **W**_SI-GEVD_ [*i*] ∈ ℝ^*N_ι_×C*^ that mapped the attended stimulus data **S**_*train*_[*i*] to the denoised EEG data 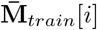 were computed using the mTRF toolbox (with the regularization parameter for ridge regularization automatically chosen (on a per trial basis) as the maximum absolute value among all the elements in the corresponding trial’s speech covariance matrix **R**_ss_). The stimulus following responses of the test trial were predicted by convolving the TRFs obtained from the training set with the stimulus envelope **S**_test_ [*i*] in the testing set:

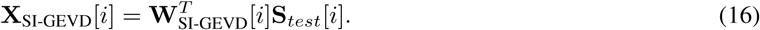

The Spearman correlation coefficients between the EEG data (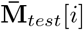 (denoised)) and the predicted stimulus following responses (**X**_SI-GEVD_[*i*]) were computed.

The correlation coefficients obtained were averaged across repetitions (from repeated stimulus trials) yielding 378 correlations (6 trials, 63 sensors). Next, we averaged the correlations in the sensor dimension using a trimmed mean (percentage = 50%) to get one correlation per trial per subject. The number of components K was chosen based on leave-one-trial-out cross-validation. For each of the 6 trials, we choose the number of components that maximized the average correlation between the predicted EEG and the actual EEG on the other 5 trials.

We benchmark the results of SI-GEVD filtering with the current state-of-the-art method for joint denoising and dimensionality reduction, namely DSS (Särelä and Valpola, 2005). It is noted that, in our case, only the isolated sentences data contains repetitions, so we can only use this part of the data for training the DSS filters. These filters then replace the SI-GEVD filters in the procedure mentioned above. The available isolated sentences repetition trials were used to extract DSS components per subject – referred to as ‘DSS-3reps’. As mentioned earlier, the SI-GEVD filters were also trained only on the isolated sentences data and thus excluding the continuous speech data in order to have a fair comparison with the ‘DSS-3reps’ case. For the normal forward model (using **W**_raw_) based on unfiltered EEG data (‘raw’), the correlations from the best 8 channels (with the highest correlations) across subjects are averaged (channels Pz, POz, P2, CPz, Oz, O2, PO3 and FC1). Note that in this context, CCA (when used as a denoising filter as in section III-B) would coincide with SI-GEVD (proof in Appendix B), hence it is not included in the analysis.

#### 2) Results

Correlations from the raw, SI-GEVD filtered and DSS-filtered data were compared using Wilcoxon’s signed-rank test with *α* = 0.05 (figure 4). The correlations from the SI-GEVD filtering data were found to be significantly higher than those of raw data (*p* = 0.0082, *W* = 81). As can be seen in figure 5, power ratio plots from DSS showed a gradual decrease of power ratio over components, indicating that the power of the stimulus following responses were spread over more than a few components, and hence choosing a small *K* would be insufficient to ensure a high SNR in the filtered data, while choosing a larger *K* will capture too much of the noise. In figure 4, it can indeed be seen that SI-GEVD filtering results in correlations significantly higher than with DSS filtering (*p* = 0.0001, *W* = 342 for ‘DSS-3reps’).

**Fig. 4:**
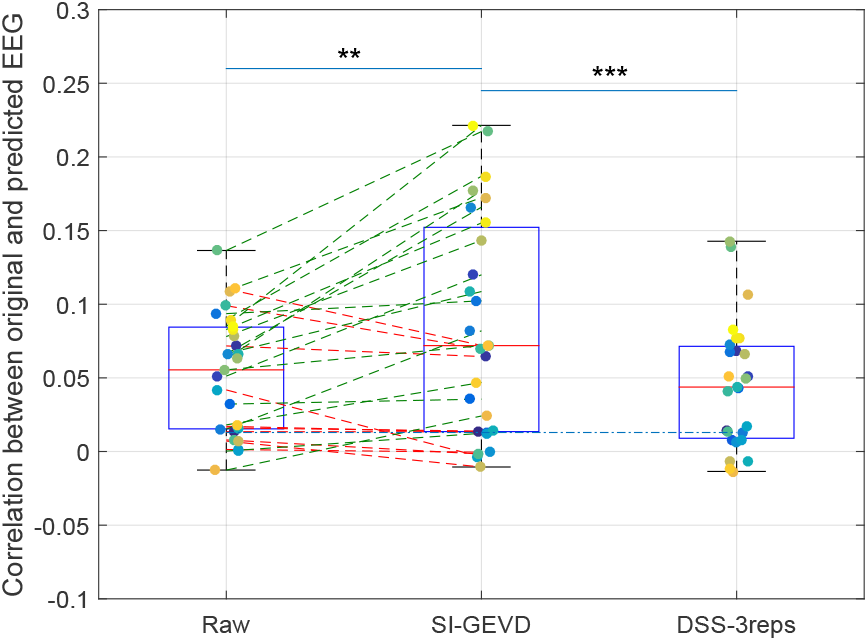
Mean correlations on 20 s trials: Comparison of mean correlations from forward modeling using raw data vs SI-GEVD filtered data vs DSS filtered data (trained only on 3 repetitions per subject (DSS-3reps)). The blue dotted line indicates the significance level (95 percentile) for performance above chance level for the correlations. Each boxplot is built from the mean correlation per subject, also indicated by colored points. Improvements in mean correlations of subjects when SI-GEVD filtering are indicated by green lines. Comparison between the methods were done using Wilcoxon’s signed-rank tests (with Bonferroni correction): ‘**’ for *p* < 0.01, ‘***’ for *p* < 0.001.

**Fig. 5:**
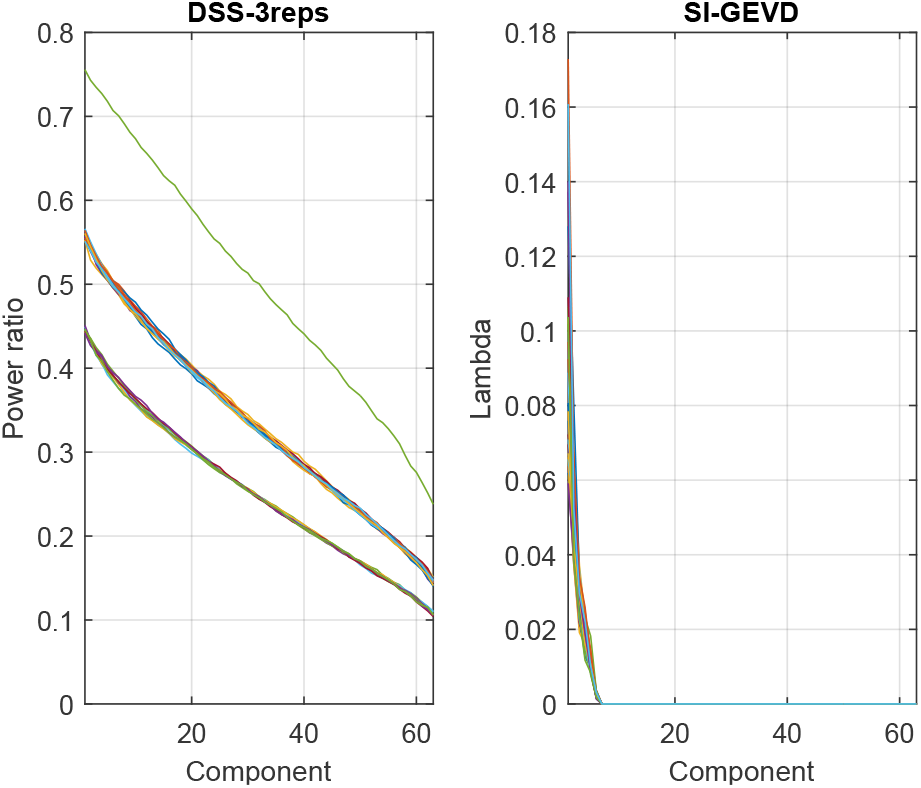
Speech envelope tracking: For the 27 subjects indicated by different colours, the above plots show the power ratios for the DSS approach and the generalized eigenvalues (λ’s) for the SI-GEVD approach (averaged across trials) for the 63 components. The drops in the curves are sharper for the SI-GEVD case showing that fewer components are necessary to ensure high SNR at the output in comparison to the DSS case.

In order to have a better understanding of the influence of the various GEVD components in the SI-GEVD approach, we investigated the weights of the back-projection matrix (each column corresponds to a component), for multiple components. Figure 6 shows the weights of each channel for the first component of the back-projection matrix (**Q**_*C*_) averaged over all training sets (each with one trial removed) and subjects. Note that the different spatial filters consist of generalized eigenvectors, which are only defined up to an arbitrary scaling and sign, which results in a corresponding inverse scaling on the columns of **Q**_*C*_. Therefore, we normalized the columns of **Q**_*C*_, so that the sum of the absolute values of the weights equal one, and corrected for sign flips before averaging the different **Q**_*C*_ matrices for each removed trial. To find those channel weights which are significantly different from zero across subjects, we used one sample cluster mass statistics (Maris and Oostenveld, 2007), with the null hypothesis that the weights of the first component of the GEVD spatial filter are symmetrically distributed around zero across subjects. A reference distribution is built by repeatedly (*n* = 5000) and randomly swapping the sign of the weights in **Q**_*K*_ across subjects and calculating a t-statistic per channel. The observed t-statistic per channel is then calculated without swapping the sign. We obtained the p-value of each channel by calculating the proportion of random samplings that have a higher t-statistic than the actual t-statistic. In this test we also take the position of the channels into account as neighbouring channels can have similar weights, using this information to reduce the family-wise error rate (details in Maris and Oostenveld (2007)). This test will show significance when the weights of the channel show a low variability and some divergence from zero over subjects. In figure 6, channels part of a cluster that is significantly different from zero (*p* < 0.01, one sample cluster mass statistics) across subjects are shown in red.

**Fig. 6:**
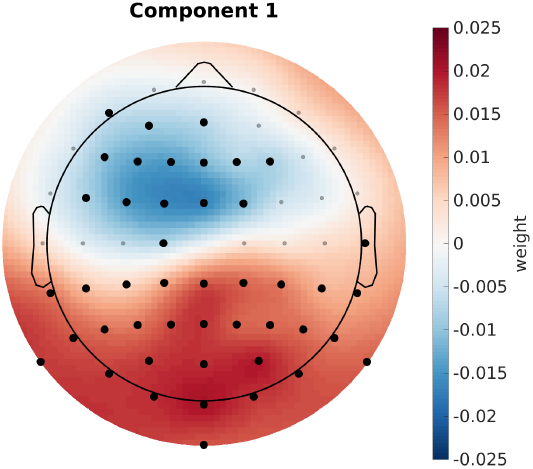
Visualization of the SI-GEVD spatial filter’s first component: Black dots indicate channels part of a cluster that is significantly different from zero.

When using only the first component, we observed two regions with an opposite polarity that contribute most to the significant difference. One is the fronto-central region, and the other is the parietal-occipital region. This is in line with other related works that estimate spatial filters based on speech processing (Dmochowski et al., 2018) and those that shows the topography of speech processing in the brain (Braiman et al., 2018; Hjortkjaer et al., 2018; O’Sullivan et al., 2015). For the second component we did not find any significant clusters. This is consistent with the results of the number of components needed to have optimal correlations and with the SNR of each component (1 GEVD component for 16 out of 27 subjects). All of this is converging evidence that the SI-GEVD filter is very effective in denoising and dimensionality reduction.

### D. Auditory attention decoding

In BCIs such as neuro-steered hearing prostheses where a person’s auditory attention, if estimated accurately, can be used to steer the noise suppression beamformer towards the direction of the attended speaker. An improvement in the SNR of the EEG data, using SI-GEVD filtering, can result in higher accuracies of AAD, resulting in better performing BCIs.

#### 1) Data Analysis

For attention decoding, we used the data from Dataset I. The preprocessed data set at a sampling rate of 32 Hz was divided into 112 trials of 20 s per subject. Adopting a leave-one-trial-out cross-validation approach, for each test trial, we used the remaining 111 trials as the training set for spatial filter estimation and TRF estimation. The difference of this application’s algorithm with that of speech envelope tracking is that here EEG is predicted from both the stimuli (attended and unattended), and an attention decoding decision is made based on how well these predictions correlate to the original EEG data, as will be explained further on. This time, we perform forward modeling in the SI-GEVD component space, and no back-projection is performed.

In the *i^th^* trial, EEG data **M**_*test*_[*i*] ∈ ℝ^*C×L*^ and the attended stimulus envelope **S**_*test*_[*i*] ∈ ℝ^*N_ι_*×*L*^ consisted of *L* = 32 Hz × 20 *s* = 640 time samples, while the training set consisted of EEG data **M**_*train*_[*i*] ∈ ℝ_*C×N–L*_ and attended stimulus envelope **S**_*train*_[*i*] ∈ ℝ^*N_ι_×N*^ with *N* =112 × *L* time samples. Stimulus envelope lags up to 400 ms were used. Therefore *N_ι_* = 32 Hz × 400 ms + 1 = 14 samples.

For each trial *i*, the training set data was used to find the SI-GEVD spatial filter **P**_*C*_ [*i*]. TRFs with respect to the attended stimulus were estimated from the training set using the SI-GEVD filtered data (no back projection). This resulted in the TRF matrix **W**_SI-GEVD_ ∈ ℝ^*N_ι_×K*^. The speech stimuli in the test trial, which was left out during the training of the TRFs and the SI-GEVD filters, was convolved with these TRFs to predict the stimulus following responses in the test trial. The predicted responses (from both the stimuli) were then compared with the SI-GEVD filtered EEG data **m**_*proj*_(*t*) by computing the Spearman correlation over each component. The stimulus that resulted in a reconstruction that yielded a higher mean correlation with the original EEG components was then considered to be the attended stimulus.

In this application, we used a different number of spatial filters *K* for each subject. An optimal number of components was found for each trial by cross-validating the difference between attended and unattended correlations when GEVD components were added one at a time, until there was no more improvement. Then, for each subject, *K* was chosen to be the highest among the optimal number of components among all trials. In the forward modeling approach using raw EEG data, the correlations from the same 8 channels as used in section III-C were averaged before making the attention decision. Similar to section III-C, we also benchmark the results against the DSS method, which is also used in a similar context for attention decoding (Akram et al., 2017). To this end, we use the trials for which the same stimulus was repeated 3 times in the EEG recording (Das et al., 2016) (4 minutes per subject). The repetition data were averaged per subject to train subject-specific DSS filters (referred to as ‘DSS-3reps’). Simulating a scenario where repetition trials are not available, we also extracted DSS components across subjects averaging trials were the same stimuli were presented to different subjects (referred to as ‘DSS-universal’). Note that such a filter training was not possible for the dataset used in speech envelope tracking (Dataset II) since the subjects were presented with different sets of stimuli.

#### 2) Results

In the SI-GEVD filtering approach, as per the criterion described before for the choice of the number of GEVD components *K*, 3 GEVD components were used for 3 subjects, 2 GEVD components for 5 subjects, and 1 GEVD component for the remaining 8 subjects. As can be seen in figure 7, the median decoding accuracy with the SI-GEVD approach was found to be 74.6%, which was 13% higher than that of raw EEG data (61.6%). Using Wilcoxon’s signed rank test, this difference was found to be significant (*p* < 0.001, *W* = 119.5). In addition, we extracted DSS components for the 2 cases: ‘DSS-3reps’ and ‘DSS-universal’. However, similar to the results from section III-C, the power ratio plots of the DSS generalized eigenvalues showed a gradual decrease of power ratio over components (see figure 8). Hence, there were not just a few dominant components that could separate the desired signal from noise. In short, we did not expect the DSS approach to meet the purpose of dimensionality reduction. As can be seen in figure 7, both the DSS approaches result in performances poorer than the SI-GEVD approach (*p* < 0.001, *W* = 134.5 for ‘DSS-3reps’ and *p* < 0.0001, *W* = 136 for ‘DSS-universal’). In the context of attention decoding, the improved SNR due to SI-GEVD filtering of the test trial can lead to higher correlations with the predicted EEG from the attended stimulus. This could be the reason for improved attended decoding results as we have seen in figure 7.

**Fig. 7:**
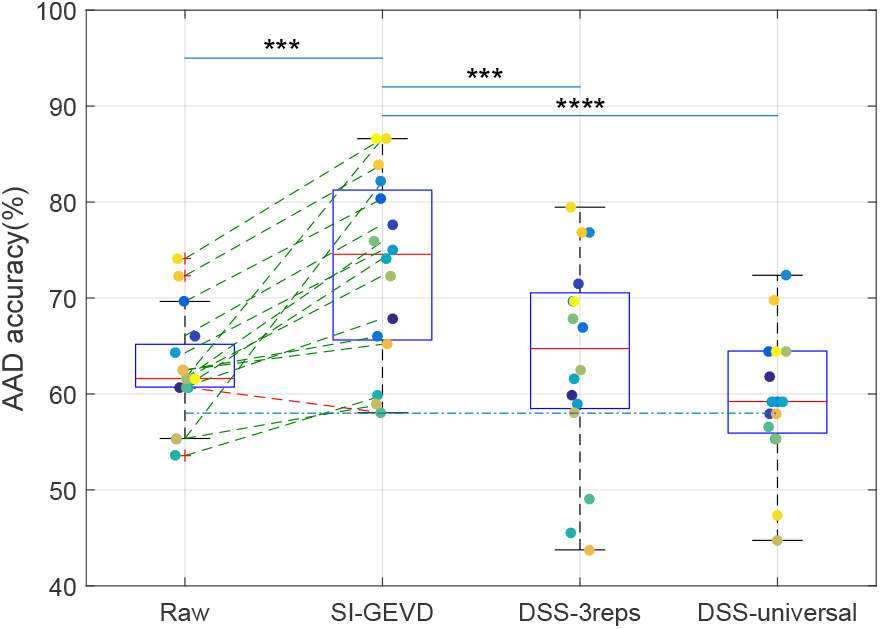
Attention decoding performance on 20 s trials: SI-GEVD filtered data vs raw data vs DSS filtered data (trained only on 3 repetitions per subject (DSS-3reps), or trained on common stimulus presentations across subjects (DSS-universal)). The blue dotted line indicates the significance level (95 percentile) for performance above chance level. Each boxplot is built from the mean decoding accuracy per subject, also indicated by colored points. Improvements in mean decoding accuracies of subjects when SI-GEVD filtering are indicated by green lines. In the plot, comparisons between methods were done using Wilcoxon’s signed-rank tests (with Bonferroni correction): ‘***’ for *p* < 0.001, ‘****’ for *p* < 0.0001.

**Fig. 8:**
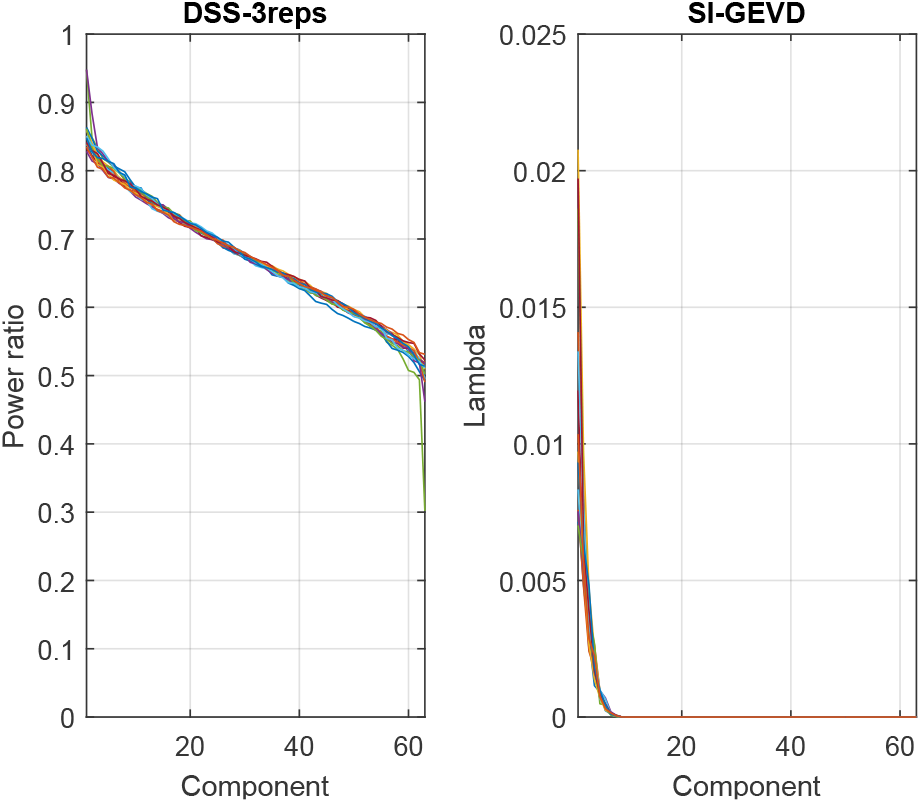
Auditory attention decoding: For the 16 subjects indicated by different colours, the above plots show the power ratios for the DSS approach (‘DSS-3reps’) and the generalized eigenvalues (λ’s) for the SI-GEVD approach (averaged across trials) for the 63 components. The drops in the curves are sharper for the SI-GEVD case showing that fewer components are necessary to ensure high SNR at the output in comparison to the DSS case.

To gain a better understanding of the denoising ability of our approach, we estimated short-term TRFs (as in section III-B) for the attended and unattended stimuli on 120 s trials, from the raw EEG as well as the SI-GEVD filtered EEG this time projected back to the electrode space and investigated whether attention decoding from the TRFs directly (as in Akram et al. (2017)) would improve. For each 120 s trial, we found the mean and standard deviation of the TRF of the channel ‘Tp7’ (one of the channels resulting in high AAD performance as shown in Narayanan and Bertrand (2019)), across trials, as shown in figure 9.

**Fig. 9:**
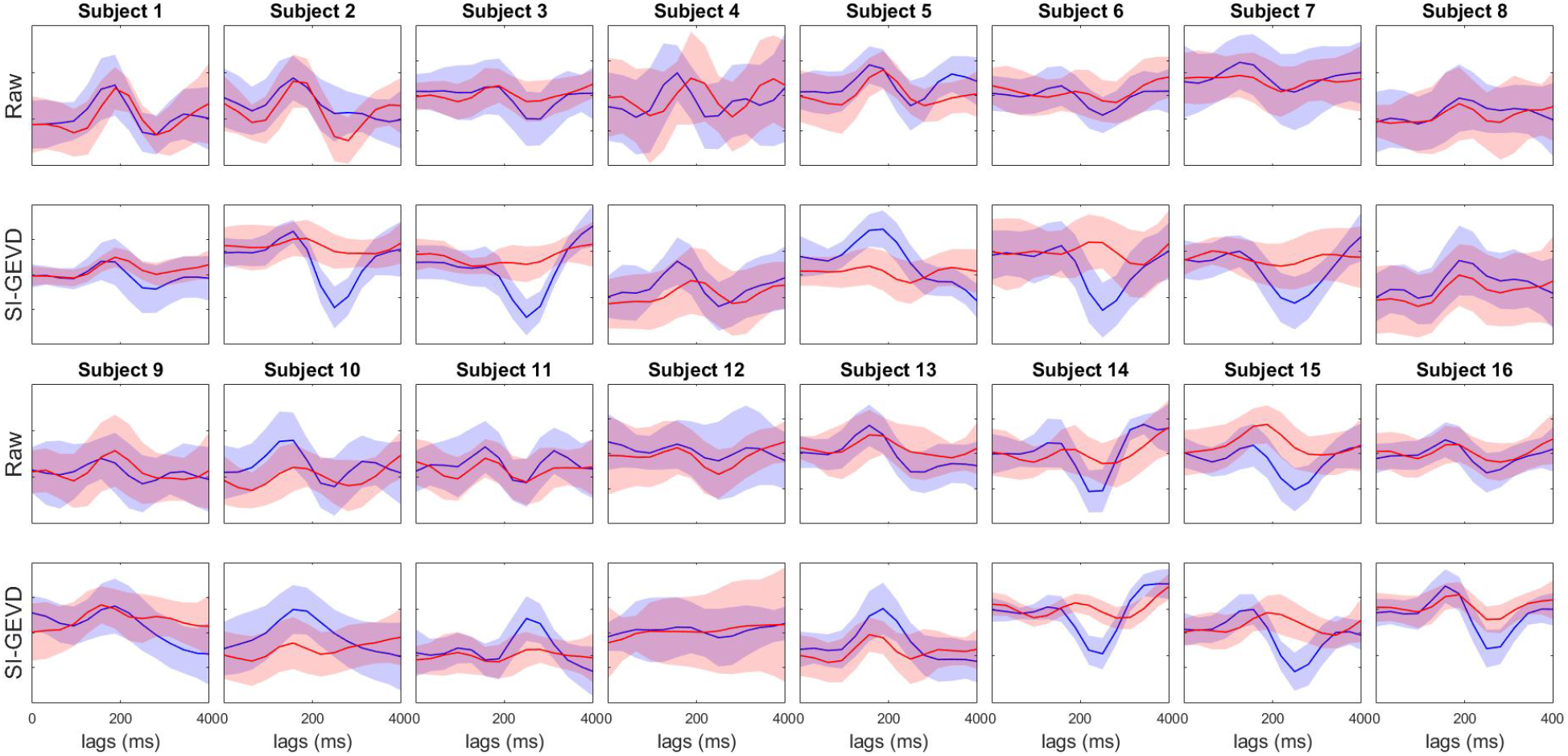
Mean attended (blue) and unattended (red) TRFs (channel ‘Tp7’, over trials) and their standard deviation (area around the mean) for the 16 subjects. The TRFs were estimated from 120 s trials of each subject. Each trial consisted of either raw EEG data (rows 1 and 3) or SI-GEVD filtering data (rows 2 and 4). The horizontal axis represents lags in ms.

We can see clear differences between the two approaches. The SI-GEVD approach results in patterns that have a better separation between attended and unattended TRFs compared to those from raw EEG data. In order to check the statistical significance, for each method we identified a reference lag at which the difference between the mean of absolute attended TRF and mean of absolute unattended TRF was maximal. We then found the number of trials for which, at the corresponding lag for each method, the absolute value of attended TRF was higher than that of the unattended TRF. Using permutation tests (detailed description in Biesmans et al. (2017)), we found that, compared to raw EEG data, SI-GEVD filtered data had significantly more trials where the absolute value (at the reference lag) of the attended TRF was higher than that of the unattended TRF (*p* < 0.0001). Knowing that peak amplitudes of short-term TRFs estimated from MEG data (which has higher SNR than EEG in general) were found to be modulated by attention (Akram et al., 2017), the better separation of attended and unattended TRFs from short-term trials of the EEG data upon SI-GEVD filtering can be seen as a result of SNR improvement.

## IV. Discussion

We presented an algorithm for joint denoising and dimensionality reduction of EEG data, in the context of auditory stimulus following responses. In order to obtain the spatial filters that project the data onto a max-SNR subspace, we employed a stimulus-informed generalized eigenvalue decomposition (SI-GEVD) of the covariance matrix of the stimulus-following neural response and the covariance matrix of the raw EEG signal. The former is estimated using a long-term forward model (TRF) between the available stimulus signal and the raw EEG data, after which the stimulus is convolved with the resulting TRF. We analyzed the performance of the proposed algorithm in 3 experiments in the context of auditory neuroscience – short-term TRF estimation, speech envelope tracking and auditory attention decoding.

In the context of TRF estimation from short trials, we used hybrid synthesized EEG in order to have access to the ground truth TRFs to assess performance. We found that, over a range of SNRs, SI-GEVD filtering effectively denoised the EEG data, resulting in TRF estimates that match better with the base TRFs, in comparison to those from EEG data without any spatial filtering. For speech envelope tracking in a single speaker scenario, the correlations between the EEG data and the predicted stimulus following responses were also found to be significantly better when using the denoised EEG data from the SI-GEVD filter, compared to DSS-filtered data or hand-picking channels. For multi-talker scenarios, SI-GEVD filtering resulted in significantly higher attention decoding accuracies than by averaging correlations over a set of raw EEG channels and then making attention decoding decisions. The attention decoding performance was also found to be better than when DSS-based spatial filtering was employed. It is important to note here that the DSS-based approach relies on data from stimulus repetitions, and the lower performance for this method can be attributed to the lack of repetitions present in our data to have a good estimate of the covariance matrix of the desired signal.

Many algorithms exist to perform dimensionality reduction and/or denoising without the use of GEVD, which we will briefly discuss below. PCA is a method that focuses on capturing most of the variance in the data without making a distinction between stimulus response and noise. This comes with the underlying assumption that the desired responses are also those that exhibit the largest variance in the data (which is definitely not the case for auditory responses). Reduced rank regression (Velu and Reinsel, 2013) is another dimensionality reduction approach where, similar to SI-GEVD, forward modeling is used to estimate stimulus-following responses, but a rank constraint is introduced on the TRF matrix. However, similar to PCA, the resulting spatial filters only aim to capture most of the variance of the stimulus following responses while ignoring the spatial characteristics of the noise. ICA is another common method for denoising, but it does not work well for extracting signal components with highly negative SNRs, i.e., in cases where the sources are weaker than the background EEG activity. ICA also does not make use of the stimulus as a side information. Another common way of dimensionality reduction is to handpick channels from brain regions that are expected to have the strongest neural responses to the stimulus of interest, or simply use all channels, and calculate a mean over these channels for the parameter (e.g., correlation) analyzed in the problem (Di Liberto et al., 2015; Wong et al., 2018; Zou et al., 2018). In section III, we have shown that this can lead to highly suboptimal results. Another method is to exhaustively train and test the system (e.g. attention decoders, or forward models) dropping one channel at a time (Fuglsang et al., 2017; Mirkovic et al., 2015; Narayanan and Bertrand, 2018) until the desired goal is achieved with fewer channels.

As explained in section II, spatial filtering to denoise neural data thereby separating stimulus-related activity from non-stimulus related activity was also addressed in de Cheveigné and Simon (2008) and de Cheveigné and Parra (2014) where the DSS procedure from Särelä and Valpola (2005) was used. DSS uses PCA to pre-whiten the data. The whitened data is then submitted to a bias function, followed by another PCA to determine the orientations that maximize the bias function, i.e., essentially the power of the biased data. The bias function averages the epochs under the same stimulus condition, thereby reinforcing stimulus-evoked activity. The second PCA step on the biased data results in a rotation matrix which can be applied to the whitened data to get signal components which can be kept or discarded depending on their bias score. The remaining components can be projected back to the sensor space resulting in denoised responses. As explained in de Cheveigné and Parra (2014), the steps of pre-whitening and PCA to find the rotation matrix can in principle be replaced by a GEVD on the correlation matrix of the biased data (with enhanced stimulus following responses) and the correlation matrix of the raw data, as done in the procedure explained in Section II. The key difference between SI-GEVD and DSS is the way we estimate the covariance matrix of the stimulus response. SI-GEVD estimates a TRF between the (known) stimulus and the recorded EEG data, after which the stimulus is convolved with the TRF to obtain an estimate of the stimulus response in each channel. de Cheveigné and Simon (2008) uses the method of epoch-averaging on pre-whitened data. Here, the stimulus does not have to be known, but a major practical limitation comes with the condition that it can only be performed if there are enough repetitions of the same stimulus in the data to effectively enhance the stimulus following responses by averaging. DSS is often used to denoise MEG responses with a relatively small number of repetitions (Akram et al., 2016; Ding and Simon, 2012b; Miran et al., 2018), but the SNR in EEG responses is considerably lower (Kong et al., 2015). In addition, the GEVD approach renders pre-whitening and the PCA steps unnecessary, resulting in improved computational efficiency. Since DSS has already been shown to outperform ICA and PCA (de Cheveigné and Simon, 2008), we did not include the latter methods in our analysis.

As opposed to PCA, ICA, DSS, JD, and many other spatial filtering techniques, the CCA method does use the stimulus signal to find relevant components in the EEG data. It does this by jointly computing the optimal projection of the stimulus and the EEG data, respectively, in order to maximize the correlation between both. We showed in Appendix B that CCA is a special instance of the proposed SI-GEVD method. However, the SI-GEVD method is more flexible to incorporate additional information such as brain function lateralization due to the direction of the stimulus. This was also experimentally verified in section III-B, where it was demonstrated that the SI-GEVD method is more effective in improving SNR compared to CCA.

The core of the SI-GEVD method is a generalized eigenvalue decomposition, which in fact plays a central role in many other EEG enhancement algorithms in the literature. However, this key role of the GEVD is not always as explicit as it was in our derivation of the SI-GEVD method. Indeed, as explained above, the JD or DSS methods in their original formulation did not involve an actual GEVD, as the equivalence with a GEVD was only identified at a later point in time. A similar example is the xDAWN algorithm (Rivet et al., 2009) which addressed the problem of dimensionality reduction in the context of brain computer interfaces – specifically a P300 speller. The goal of discriminating epochs containing a P300 potential was tackled by estimating the P300 subspace from raw EEG data, and projecting the raw EEG onto this subspace, effectively enhancing the P300 evoked potentials (Rivet et al., 2011, 2009). The xDAWN algorithm uses either an averaging operation across time-locked responses (similar to DSS) or a forward encoding model (similar to SI-GEVD) where the stimulus onset triggers are used as the regressor to estimate a template for the event-related potentials (ERPs), and consequently, the second order statistics of the neural responses evoked by the target stimulus pulses. In Rivet et al. (2011), it has been identified that the two core algebraic operations in xDAWN (the QR and singular value decomposition) used for estimation of the spatial filter can be replaced by a single GEVD on the matrix pair of the correlation matrices of the stimulus evoked responses and the raw EEG data. As such, xDAWN can be viewed as a special case of DSS or of SI-GEVD, albeit for the case of pulsed stimuli, where the analysis is driven by time triggers rather than continuous stimulus waveforms as targeted in this paper. It can be easily shown that a time-locked averaging becomes equivalent to a forward regression based on time triggers when the time between any two consecutive stimulus pulses is larger than the neural response to each of them. Therefore, in this particular case, xDAWN, DSS, SI-GEVD, and consequently also CCA (see Appendix B), all collapse to one and the same procedure, which reveals an interesting link between these -at first sight-very different methods.

Another algorithm that relies on GEVD is the common spatial pattern analysis (CSP) (Dornhege et al., 2006; Müller-Gerking et al., 1999; Ramoser et al., 2000) which aims to solve a binary classification task. To achieve this, the algorithm finds a set of spatial filters that maximize the variance of the projected data for one class, and minimize the variance for the other, such that the output power of the filters can be used as features in a classification task. The solution is found by a GEVD, yet with the main purpose to maximize the discriminative power of the output features, rather than maximizing SNR. Therefore, the GEVD is applied to the two covariance matrices corresponding to both classes, which is akin to Fisher’s linear discriminant analysis. Similar to the DSS approach in de Cheveigné and Simon (2008), in CSP, epochs of data are sometimes averaged, per class, in order to have a good estimate of the covariance matrix of the desired signal. Source power comodulation (SPoC) (Dähne et al., 2014) is another GEVD-based algorithm that finds a spatial filter to decompose the neural data into components that are maximally correlated with the intensity of the auditory input. The goal is then to maximize the co-modulation between the power time course of the spatially filtered signal and the target variable (stimulus intensity). These algorithms, even though they employ GEVD, are different from SI-GEVD in terms of the underlying problem they try to solve and the definition/calculation of the (covariance) matrices that are used in the GEVD.

## V. Conclusion

Dimensionality reduction and denoising are important steps in the procedure to analyze neural responses, and particularly stimulus-following responses in EEG data which often has low SNRs. In this context, methods like DSS and PCA are often used in order to denoise and/or reduce the dimensionality of the data. When it comes to forward modeling approaches, often researchers handpick channels based on their knowledge of locations of the strongest stimulus following cortical activity. In this paper, we present and evaluate a stimulus-informed GEVD-based filtering approach which makes use of predicted stimulus following responses to find a spatial filter that maximizes the SNR at its output. Our method is fully data-driven, and has the advantage of not relying on repeated trials for the spatial filter estimation. We have shown the benefits of using our approach by analyzing 3 different applications in the field of auditory neuroscience, and compared the performance of the proposed method with other state-of-the-art approaches like CCA, DSS or averaging over selected channels.

## Appendix

### A. Optimization problem equivalence

The SI-GEVD filter is found by solving of the optimization problem

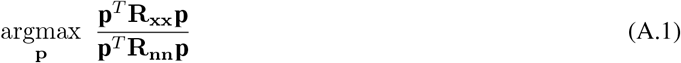

resulting in the GEVD of the matrix pencil (**R_xx_**, **R_nn_**). It is shown below that solving the optimization problem for maximizing the SSNR (derivation was taken from Wouters et al. (2018))

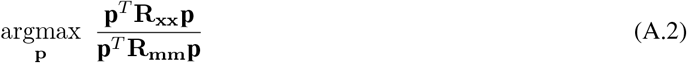

results in the same solution as that for maximizing the SNR (A.1).

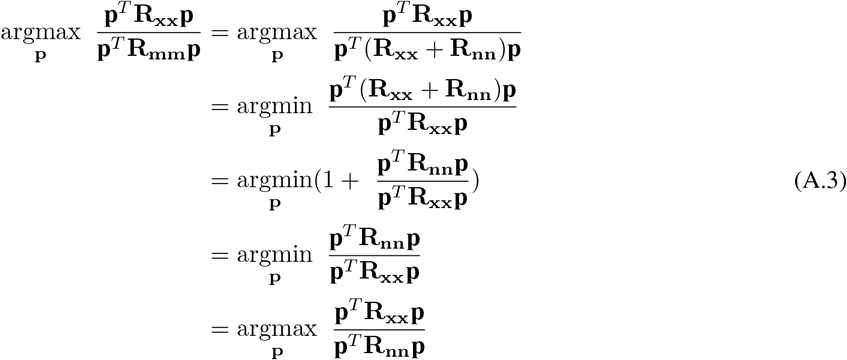

Thus, it can be concluded that the GEVD of the matrix pencil (**R_xx_**, **R_mm_**) would result in the same SI-GEVD filter as for (**R_xx_**, **R_nn_**).

### B. Equivalence to canonical correlation analysis (CCA)

The SI-GEVD spatial filter p can be found by solving

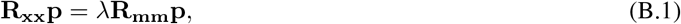

where the equivalence of the solution to (9) has been shown in Appendix A. From equations (4),(6) and (7), the stimulus following response **X** can be estimated as

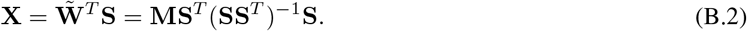

In the SI-GEVD procedure, the spatial covariance matrix of the stimulus following response **X** is estimated as **R_xx_** ≈ (**XX**^*T*^)/*N*. Using equation (B.2), we have

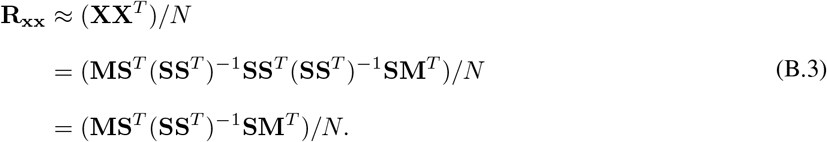

In addition, the spatial covariance matrix of the raw EEG data **R_mm_** is estimated as **R_mm_** ≈ (**MM**^*T*^)/*N*. The equation (B.1) can, therefore, be rewritten as

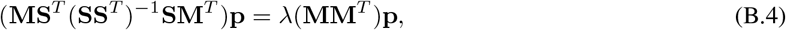

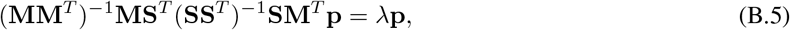

where the division my *N* has been removed from both sides. The SI-GEVD filter can thus be found as the eigenvector corresponding to the maximum eigenvalue in the eigenvalue decomposition of (**MM**^*T*^)^−1^**MS**^*T*^(**SS**^*T*^)^−1^**SM**^*T*^.

Now let us consider using canonical correlation analysis (CCA) to find a spatial filter **p_m_** ∈ ℝ^*C*^ for the raw EEG data m(t) and a temporal filter **p_s_** ∈ ℝ^*N_ι_*^ for the stimulus envelope (including *N_ι_* – 1 lagged versions of it) *s*(*t*), such that the spatially filtered neural response **m**_*cca*_(*t*) = **p_m_**^*T*^**m**(*t*) and the temporally filtered stimulus **s**_*cca*_(*t*) = **p_s_**^*T*^**s**(*t*) are maximally correlated. The optimization problem maximizing this correlation is given by (Friman et al., 2002)

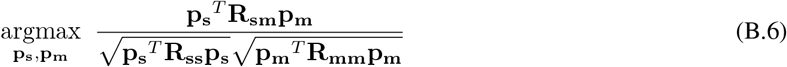

where cross-correlation matrix **R_sm_** = *E*{**s**(*t*)**m**(*t*)^*T*^} ∈ ℝ^*N_ι_×C*^ can be estimated as (**SM**^*T*^)/*N*, and the EEG and stimulus envelope covariance matrices can be estimated as **R_mm_** ≈ (**MM**^*T*^)/*N* ∈ ℝ^*C×C*^ and *R_ss_* ≈ (**SS**^*T*^)/*N* ∈ ℝ^*N_ι_×N_ι_*^ respectively.

Solving this optimization problem, spatial filter **p_m_** for the EEG data is obtained as the eigenvector corresponding to the highest eigenvalue of the eigenvalue problem (Friman et al., 2002; Hotelling, 1936):

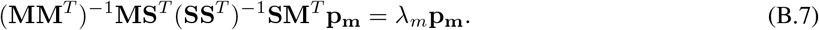

Remarkably, when comparing this equation (B.5), we find that the SI-GEVD algorithm, which aims to maximize the SNR of the EEG data (where the desired signal is defined as the stimulus following response) and the CCA algorithm, which aims to maximize the correlation between the EEG data and the stimulus by means of a joint transformation on both modalities result in the exact same solution. This proves that, in its simplest form, the SI-GEVD boils down to CCA, making the latter a special case of the former. It is noted that this equivalence breaks down if temporal information is also incorporated on the EEG side through the use of time lags (similar to what is done now on the stimulus side).

### C. Back-projection to the electrode space

If **P_C_** contains the set of generalized eigenvectors of the matrix pencil (**R_xx_**, **R_mm_**) in its columns, and 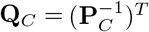, it follows that

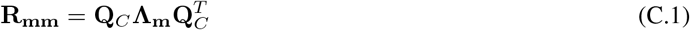

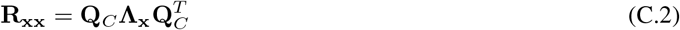

where **Λ_m_** and **Λ_x_** are diagonal matrices such that 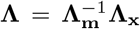 (Golub and Van Loan, 1996). Here **Λ** is the diagonal matrix containing all the generalized eigenvalues. The goal is to find a filter **V** ∈ ℝ^*K×C*^ which projects the compressed *K*-component data **m**_*proj*_(*t*) back to the electrode space with minimal error in the least squares sense:

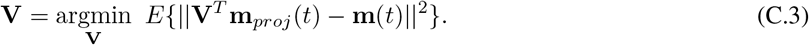

This is a standard minimum mean squared error problem of which the solution is given as

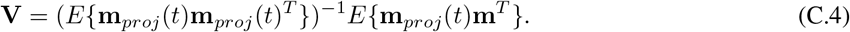

This can be rewritten using the notation introduced in section II as

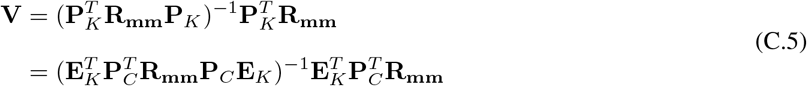

where **E**_*K*_ ∈ ℝ^*C×K*^ is a matrix that chooses the first *K* columns of **P**_*C*_ so that **P**_*K*_ = **P**_*C*_**E**_*K*_. Since 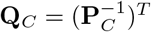, from (C.1) it follows:

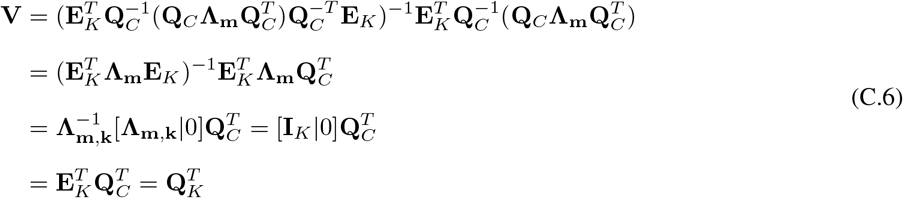

where **Λ_m,k_** is a diagonal matrix containing the first *K* diagonal values of **Λ_m_**, and [**Λ_m,k_**|0] represents the concatenation of **Λ_m,k_** with *C* – *K* columns of all zero values (equivalent to choosing the first *K* rows of **Λ_m_**). The derivation shows that the filter that should be used to back-project **m**_*proj*_(*t*) to the electrode space is 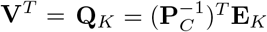.

## Acknowledgments

The authors would like to thank all the subjects for their participation in the study.

## Footnotes

1 In some applications, one may want to work on the compressed EEG signals directly rather than reconstructing the neural response in the electrode space. For example, in Akram et al. (2017) short-term TRFs are estimated on a single component in the compressed space to track an M100 peak.

2 Cz as the reference was an arbitrary choice. It is commonly used since electrodes in the ‘Z’ sites won’t necessarily reflect or amplify lateral hemispheric cortical activity as they are placed over the corpus callosum.

3 The back-projection is based on a least squares estimator, which is known to be biased in the case of noise in the regressor matrix.

